# Transcranial Direct Current Stimulation Modulates Resting Brain Hemodynamics and Autonomic Function: A Multimodal fNIRS-HRV Study

**DOI:** 10.1101/2025.09.07.673917

**Authors:** Seoyoung Choi, Sungmin Ha, Woong Park

## Abstract

Transcranial direct current stimulation (tDCS) is a non-invasive neuromodulation technique that can influence brain activity and physiological function. We conducted a sham-controlled multimodal study to examine the effects of low-intensity (0.375 mA) bifrontal tDCS at rest in healthy adult males. The tDCS, with the anode placed over the left dorsolateral prefrontal cortex (DLPFC) and the cathode over the right DLPFC, was applied for twelve minutes, and outcomes were measured with functional near-infrared spectroscopy (fNIRS) for cortical hemodynamics and connectivity, heart rate variability (HRV), photoplethysmography (PPG) for autonomic function, and subjective surveys for emotional stress. The results showed that, compared to sham, active tDCS produced a decrease in oxyhemoglobin (HbO) concentration in the left DLPFC during stimulation, indicating reduced cortical oxygenation in the stimulated region. Functional connectivity analysis of fNIRS signals further revealed altered network connectivity, including modulation of intra- and inter-frontal connections, in the tDCS condition relative to sham. Concurrently, tDCS induced an increase in HRV indices such as RMSSD, SDNN, pNN50, and Poincaré plot measures SD1 and SD2, reflecting enhanced parasympathetic activity and autonomic regulation. Participants in the active tDCS group also reported selective reductions in both state and trait anger after stimulation, whereas the sham group showed minimal changes in mood. These findings demonstrate that even at a subthreshold intensity, bifrontal tDCS at rest engaged neurovascular and neurovisceral mechanisms, coupling changes in prefrontal cortical activity with autonomic outflow and mood. This study provides new evidence of cortical-autonomic coupling during neuromodulation and suggests that low-intensity frontal tDCS may promote a calm physiological and emotional state. The results have implications for future research on brain stimulation in emotion regulation and for developing neuromodulation-based interventions to improve autonomic balance and mood in both healthy individuals and clinical populations.

## 1. Introduction

Transcranial direct current stimulation (tDCS) is a form of non-invasive brain stimulation that applies a weak direct current (typically 0.5–2 mA) to the scalp to subtly modulate neuronal membrane potentials (Nitsche & Paulus, 2000; Reed, & Cohen, 2018; Santarnecchi et al, 2015). By placing anodal and cathodal electrodes over target regions, tDCS can induce polarity-specific changes in cortical excitability: anodal stimulation generally depolarizes neurons (increasing excitability), whereas cathodal hyperpolarizes them (decreasing excitability) (Bouchard et al., 2023). The dorsolateral prefrontal cortex (DLPFC) is a frequent target for tDCS due to its involvement in cognitive control, emotion regulation, and its widespread connections to other brain networks. Bifrontal DLPFC montages (with anode over left DLPFC and cathode over right DLPFC) have been used in neuromodulation studies and clinical trials (e.g. for depression) to augment left-prefrontal activity and behavioral outcomes (Brunoni et al., 2011; Carnevali et al., 2020). However, the physiological effects of low-intensity tDCS – at currents well below 1 mA – are not well characterized. In the present study, we applied a very low current (0.375 mA, significantly below the conventional range) in a bifrontal montage to explore whether such gentle stimulation can produce measurable changes in the brain and body at rest. Using a multimodal approach, we examined cortical hemodynamics, functional connectivity, autonomic nervous system activity, and mood in order to capture a comprehensive picture of tDCS effects.

Combining tDCS with functional neuroimaging and physiological monitoring can reveal how modulating the cortex influences broader neurovascular and neurovisceral systems (Nikolin et al., 2017). Functional near-infrared spectroscopy (fNIRS) is well-suited for concurrently measuring cortical blood oxygenation changes during tDCS, providing an index of regional neural activity and neurovascular coupling. Prior studies using fNIRS with tDCS have shown that tDCS can indeed alter cerebral perfusion and oxygenation signals (Bouchard et al., 2023; Stagg et al., 2013). For example, a systematic review reported that anodal tDCS often leads to detectable changes in fNIRS signals at the stimulation site (Figeys et al., 2021). At-rest recordings have sometimes found increased cortical activation (higher oxyhemoglobin) under the anode (Patel et al., 2020), although findings vary with stimulation parameters and baseline brain states. In addition to local hemodynamics, tDCS may influence functional network connectivity. The DLPFC is a hub within frontoparietal and limbic networks, so modulating this region’s excitability could reconfigure network interactions. Studies using resting-state fMRI and fNIRS have noted that prefrontal tDCS can change connectivity patterns across distributed networks (Ishihara et al, 2025; Keeser et al, 2011; Patel et al., 2020; Yaqub et al., 2018). For instance, anodal DLPFC tDCS has been found to reduce excessive intrahemispheric connectivity in the left prefrontal cortex, suggesting a focal reorganization of network dynamics (Ishihara et al, 2025). We aimed to extend these findings by assessing functional connectivity changes via fNIRS, hypothesizing that low-intensity tDCS would induce subtle but meaningful network modulations even at rest.

In parallel, we investigated autonomic nervous system responses to tDCS by measuring heart rate variability (HRV) and peripheral pulse via photoplethysmography. The prefrontal cortex is a key node in the central autonomic network and is known to modulate autonomic outflow (Beissner et al., 2013; Gomez-Alvaro et al., 2024; Thayer & Lane, 2009; Thayer et al., 2012). High prefrontal activity is associated with greater parasympathetic (vagal) tone and inhibitory control over stress responses, according to the neurovisceral integration model (Nikolin et al., 2017). Conversely, impaired prefrontal regulation is linked to autonomic dysregulation in conditions like stress and chronic pain (Gomez-Alvaro et al., 2024). There is growing evidence that prefrontal tDCS can influence HRV, presumably by engaging cortico-vagal pathways. For example, anodal stimulation over DLPFC has been shown to increase high-frequency HRV components (reflecting vagal activity) in healthy individuals (Nikolin et al., 2017). A recent randomized trial reported that anodal left DLPFC tDCS at 1–2 mA significantly increased indices such as HF power and RMSSD (root-mean-square of successive beat interval differences) compared to sham (Gomez-Alvaro et al., 2024). These changes indicate a shift toward parasympathetic dominance. We therefore expected that even low-intensity bifrontal tDCS might elicit detectable autonomic effects, such as an increase in HRV and a reduction in heart rate, consistent with enhanced parasympathetic modulation (Gu et al., 2022).

Beyond separate measures of brain and heart activity, this study incorporates a novel index of cardio-cerebral coupling. We compute the phase locking value (PLV) between the peripheral pulse (photoplethysmogram, PPG) and fNIRS oscillations. Low-frequency oscillations (∼0.1 Hz) in blood volume and oxygenation are known to be present in fNIRS/BOLD signals at rest, partly driven by systemic rhythms like blood pressure Mayer waves and respiration (Obrig et al., 2000; Tong & Frederick, 2010). In fact, a substantial fraction of “resting-state” hemodynamic fluctuations can originate from these systemic sources (Tong et al., 2019). By quantifying the phase synchronization between the cardiac pulse and cerebral hemodynamic signals, we obtain insight into the degree of heart-brain physiological coupling. Changes in this coupling might indicate alterations in cerebral autoregulation or vascular tone influenced by ANS activity. We ask whether tDCS modulates this heart-brain interaction – for example, if tDCS increases parasympathetic activity, it could affect cerebrovascular oscillatory patterns or timing, thus changing the PLV.

Finally, we included subjective psychological assessments to probe whether physiological changes with tDCS correspond to any acute mood or stress alterations. The prefrontal cortex plays a prominent role in emotion regulation, and altering its activity can impact mood and affective states (Abend et al., 2019; Liu et al., 2017; Morgan et al., 2014). We focused on measures of emotional stress (both state – the current feeling – and trait – a disposition) because DLPFC-mediated executive control is thought to help suppress emotion like anxiety, anger and aggressive impulses. Some prior work has explored tDCS as a tool to modulate aggression and anger. For instance, stimulation of ventromedial PFC has been associated with reduced aggressive behavior in provocation tasks (Summerell et al., 2024). However, a recent meta-analysis of tDCS effects on anger and aggression found no significant overall effect across studies (Denson et al., 2025), possibly due to heterogeneity in methods and underpowered samples. Given this mixed evidence, it remains valuable to examine mood-related outcomes in new tDCS paradigms. We reasoned that low-level DLPFC stimulation during rest might exert a mild calming effect, reflected in reduced self-reported stress, consistent with the notion that enhancing prefrontal control can suppress negative affect. Such an outcome would underscore the functional relevance of cortical-autonomic coupling for emotional state regulation.

In summary, this study was designed to investigate the multifaceted impacts of low-intensity bifrontal tDCS at rest on: (1) cortical hemodynamics in the DLPFC (via fNIRS oxyhemoglobin changes) and prefrontal functional connectivity, (2) autonomic nervous system activity (HRV and pulse metrics), (3) PLV to assess heart-brain physiological coupling, and (4) emotional stress. Healthy adult males were tested to minimize hormonal variability and because men’s emotional responses have been relatively understudied in tDCS research. We hypothesized that relative to sham, active tDCS would modulate the fNIRS signals over the DLPFC (reflecting changes in cortical activation and blood flow), alter the connectivity between prefrontal regions, increase HRV (indicating enhanced parasympathetic tone and calmer autonomic state), and reduce emotional stress levels. By integrating these measures, we aimed to shed light on the neurovascular and neurovisceral mechanisms through which even a weak direct current can influence brain function and bodily state. Demonstrating effects with only 0.375 mA of current would also have practical significance, suggesting that ultra-low-dose neuromodulation can engage brain networks and potentially yield beneficial physiological and emotional outcomes with minimal stimulation intensity.

## 2. Methods

### 2.1. Participants

Thirty-five healthy male undergraduate students from Yonsei University (aged 19–28 years) participated in this study after providing written informed consent. Participants were randomly assigned to either the active tDCS group (n = 24) or the sham control group (n = 11). All individuals were screened to ensure no history of neurological, cardiovascular, or psychiatric conditions, and no contraindications to non-invasive brain stimulation. The study protocol was approved by the Institutional Review Board of Yonsei University (IRB No. 7001988-202408-HR-2369-01).

A total of 32 participants (tDCS = 22, sham = 10) were included in the final analysis due to data collection errors and low signal quality (e.g., fNIRS channel rejection, HRV artifacts), with valid datasets differing slightly by modality.

### 2.2. Experimental Design and Procedure

We employed a between-subjects, sham-controlled design with repeated measures across three blocks: pre-stimulation, during-stimulation, and post-stimulation. Participants arrived at the laboratory and were briefed on the procedure before providing written informed consent. Prior to equipment setup, they completed a battery of self-report questionnaires, including the emotional stress and items assessing physical and emotional health status. Following this, the experimenter assisted each participant in donning the equipment, including fNIRS, tDCS, and PPG sensors. Once all devices were calibrated, the resting-state recording began. Throughout the session, participants were instructed to relax, remain still, and focus on a fixation cross presented at the center of a monitor. Upon completion, the devices were removed, and participants completed post-stimulation questionnaires, including the same stress and health measures as well as a brief tDCS safety and comfort assessment.

Participants were randomly assigned to either the active tDCS group or the sham control group. The resting-state period was divided into three sequential blocks: a 6-minute pre-stimulation baseline (prestim), a 12-minute stimulation period (durstim), and a 12-minute post-stimulation period (poststim).

For the tDCS group, a constant current stimulation was delivered during the durstim block using a neuroConn DC-Stimulator Plus device (neuroConn, Ilmenau, Germany). The stimulation intensity was set to 0.375 mA—a low amplitude chosen to ensure safe current density given the relatively small 3 x 3 cm (9 cm^2^) electrode size, while still engaging cortical circuits (0.042 mA/cm^2^) (Thair et al., 2017). The anodal electrode (custom-made 3 × 3 cm hydrogel patch with adhesive and conductive properties) was placed over the left dorsolateral prefrontal cortex (DLPFC), corresponding to the F3 position of the 10–20 EEG system. The cathodal electrode (3 × 3 cm) was placed over the right DLPFC (F4 position). Stimulation was applied for the full duration of the 12-minute durstim block.

In the sham group, electrodes were placed in the same positions, but no current was delivered (single-blind design). Participants were blinded to their group assignment. The poststim block immediately followed the stimulation or sham period to assess potential after-effects of neuromodulation. The total session, including preparation, equipment setup, and brief inter-block transitions, lasted approximately 40 to 45 minutes.

### 2.3. Measures and Data Analysis

#### 2.3.1. fNIRS Recording

Cerebral hemodynamic activity was recorded using a continuous-wave functional near-infrared spectroscopy (fNIRS) system, NIRSIT Lite (OBELAB Inc., Seoul, Korea). This portable and wireless device employs dual-wavelength near-infrared light (approximately 780 nm and 850 nm) to measure changes in oxygenated (HbO) and deoxygenated (HbR) hemoglobin concentrations in cortical tissue. The headset contains 5 light sources and 7 detectors, yielding a total of 15 measurement channels that are symmetrically distributed across the prefrontal cortex. The source-detector separation is optimized (∼30 mm) for cortical sensitivity, and the data were sampled at 8.138 Hz throughout the resting-state recording.

Before the experiment, the device was positioned over the participant’s forehead, covering the bilateral dorsolateral and frontopolar prefrontal cortex, as illustrated in Fig. 1. Signal quality was checked and calibrated prior to recording to ensure low optical noise and sufficient gain across all channels. During the resting-state session, raw optical intensity data were continuously recorded, and event markers were logged to segment the data into prestim, durstim, and poststim blocks. Hemodynamic changes were calculated using the modified Beer–Lambert Law (MBLL), with adult-specific differential pathlength factors (DPFs) applied. We focused primarily on changes in oxygenated hemoglobin (HbO), which are more sensitive to regional cerebral blood flow and neural activation.

**Figure 1.**
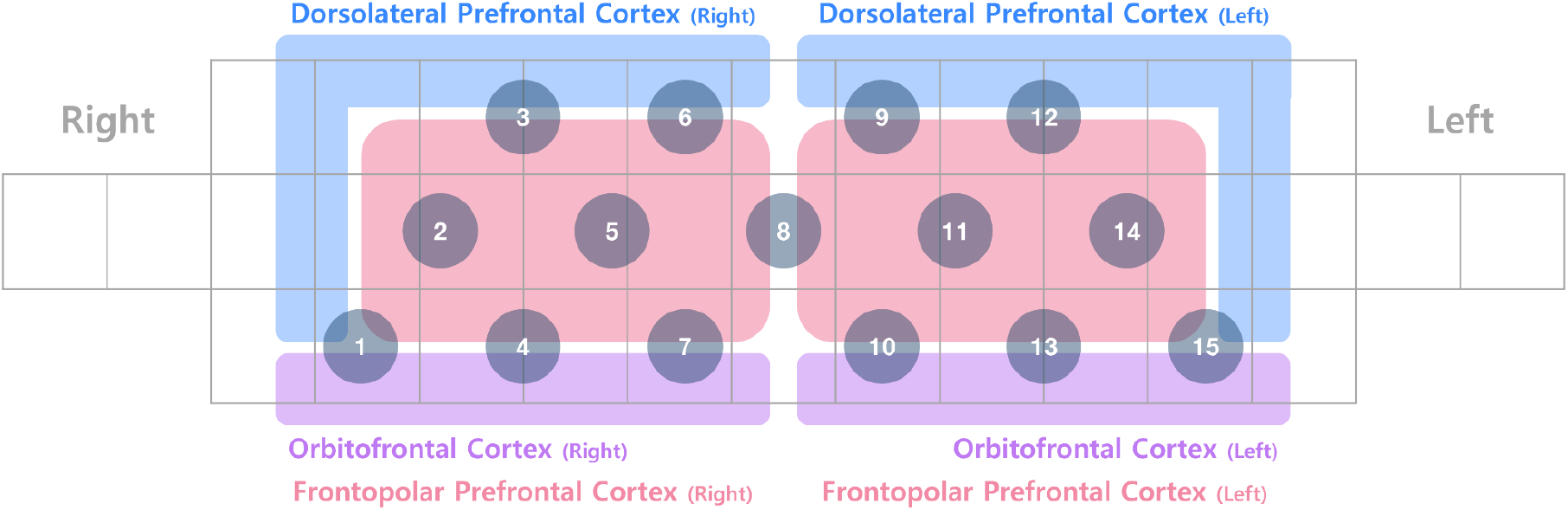
Channel locations of the NIRSIT Lite fNIRS device. Fifteen channels are positioned over the prefrontal cortex, covering regions within the anterior prefrontal cortex, including the bilateral dorsolateral prefrontal cortex (DLPFC), orbitofrontal cortex(OFC), and frontopolar cortex (FPC).

Preprocessing of the fNIRS data was performed using NIRSIT QUEST v1.0.4, a software platform developed by OBELAB and based on MATLAB (MathWorks, Natick, MA). The preprocessing pipeline included several steps to ensure high signal quality and physiological interpretability of the fNIRS signals. First, channels with invalid or missing raw light intensity values were excluded. Additional channel rejection criteria were applied based on (1) low median intensity values (<30 a.u.) (Yücel et al., 2021), (2) high temporal variability (coefficients of variation >7.5% in more than 10% of 5-second windows) (Bonilauri et al., 2021), and (3) prolonged signal saturation (i.e., consecutive identical values exceeding 5% of the recording). After artifact-prone channels were removed, raw intensity data were converted to optical density (OD) (Yücel et al., 2021), and motion artifacts were corrected using the Temporal Derivative Distribution Repair (TDDR) algorithm (Fishburn et al., 2019). OD data were then converted into hemoglobin concentration changes using the modified Beer–Lambert law (Delpy et al., 1988; Cope, & Delpy, 1988), with differential pathlength factors (DPFs) estimated based on participant age (Scholkmann, & Wolf, 2013), and molar extinction coefficients calculated by Moaveni (Zhao et al., 2017). The resulting concentration data were expressed in micromolar (μM) units. To remove slow drifts and high-frequency noise, a 4th-order Butterworth band-pass filter (0.005–0.1 Hz) was applied to the hemoglobin signals (Yücel et al., 2021). Channels showing extremely negative correlation (*r* < –0.9) between HbO and HbR were discarded as physiologically implausible (Takizawa et al., 2014). Finally, remaining motion artifacts were attenuated using Correlation-Based Signal Improvement (CBSI), which leverages the expected inverse relationship between HbO and HbR during true neural activation (Cui et al., 2010). For subsequent statistical analyses, we focused primarily on oxygenated hemoglobin (HbO) signals due to their higher sensitivity to changes in regional cerebral blood flow and neural activation.

#### 2.3.2. Heart Rate Variability (HRV)

A finger photoplethysmography (PPG) sensor was attached to the index finger of the participant’s non-dominant hand to record the pulsatile blood volume waveform continuously at a sampling rate of 100 Hz. PPG data were segmented into three experimental blocks - prestim: 6 min; durstim: 12 min; poststim: 12 min - according to task timing. Raw signals were preprocessed using NeuroKit2’s function to reduce motion artifacts and baseline drift. Cardiac pulses were detected with a minimum inter-peak distance of 0.4 s (distance = 0.4 × sampling rate) and a prominence threshold of 0.5. Inter-beat intervals (IBIs, in milliseconds) were computed from successive peak onsets, and physiologically implausible intervals (< 350 ms or > 2000 ms) were excluded from analysis. As a result, the final HRV sample was 12 participants in the tDCS group and 6 in the sham group.

HRV indices were computed using NeuroKit2’s hrv function (sampling rate = 100 Hz). Time-domain metrics included SDNN, RMSSD, and pNN50; frequency-domain metrics included LF, HF, and LFHF; nonlinear metrics from Poincaré plot analysis included SD1 and SD2; and additional nonlinear measures included approximate entropy (ApEn) and detrended fluctuation analysis (DFA-α_1_). All indices were computed for each block and participant using NeuroKit2’s validated pipeline (Makowski et al., 2021). Mean heart rate (beats per minute) was calculated for each block by dividing the number of detected peaks by block duration in minutes.

Each HRV metric was interpreted according to established physiological meaning. SDNN reflects overall HRV and total variability across recording periods. RMSSD and pNN50 primarily index high-frequency variability mediated by parasympathetic activity (Laborde et al., 2017). LF is associated with both sympathetic and parasympathetic influences, whereas HF reflects parasympathetic modulation linked to respiratory sinus arrhythmia. The LF/HF ratio has been used as a marker of sympathovagal balance. SD1 and SD2 represent short- and long-term variability, respectively, in Poincaré plot geometry. ApEn quantifies signal complexity and regularity, and DFA-α_1_ captures short-term fractal-like correlations in the IBI series.

Statistical analyses were performed in RStudio (version 4.3.3). Each HRV metric and mean heart rate was analyzed using a 2 (Group: tDCS, sham) × 3 (Block: prestim, durstim, poststim) repeated-measures ANOVA. Significant effects were followed by Bonferroni-adjusted pairwise comparisons. Additionally, within-group one-way repeated-measures ANOVAs across blocks were conducted, with Bonferroni-corrected paired *t*-tests for follow-up comparisons, and between-group *t*-tests were performed at each Block. Normality was assessed using Shapiro–Wilk tests, and skewed variables (e.g., pNN50) were log- or square-root-transformed as needed prior to ANOVA.

#### 2.3.3. Phase Locking Value (PLV) between PPG and fNIRS

To quantify cardio–cerebral coupling, we computed the phase-locking value (PLV) between the slow-band fingertip PPG envelope and prefrontal oxyhemoglobin (HbO) measured by fNIRS. For each participant, HbO time series from the 15 main channels (Channels 1–15; native sampling 8.138 Hz) and the PPG waveform (100 Hz) were preprocessed as follows: the PPG was band-pass filtered in the cardiac range (0.7–8 Hz) and its analytic amplitude (envelope) was obtained via the Hilbert transform; both the PPG envelope and HbO were then band-pass filtered in the 0.07–0.15 Hz range using zero-phase Butterworth filters, and the slow-band PPG envelope was linearly aligned to the fNIRS time base. Instantaneous phases were extracted with the Hilbert transform.

PLV was computed for each channel and experimental block (prestim: 0–360 s; durstim: 360–1080 s; poststim: 1080–1800 s) as the modulus of the time-average of unit phasors formed from the phase difference between HbO and PPG, yielding values in [0, 1] with higher values indicating stronger phase synchronization (Lachaux et al., 1999). For statistical analysis at the region-of-interest (ROI) level, channel-wise PLVs (Channels 1–15) were averaged within each block and participant.

All preprocessing and analysis were performed in Python 3 using the SciPy library (Virtanen et al., 2020) for filtering and tests; the statsmodels (Seabold & Perktold, 2010) and/or pingouin for mixed ANOVA. Multiple comparisons in channel-wise tests were addressed with FDR correction (Benjamini–Hochberg). Significance was set at α=.05 (two-tailed).

#### 2.3.4. Subjective Surveys

To assess participants’ perceived emotional stress, we administered the Emotional Stress Inventory (ESI) (Chon et al., 2020) immediately before the prestim block and immediately after the poststim block. This self-report scale consists of 42 items covering a range of state anger, state anxiety, state depression, trait anger, trait anxiety, and trait depression, rated on a 6-point Likert scale (1 = “strongly disagree” to 6 = “strongly agree”). The inventory encompasses multiple domains such as tension, worry, sensitivity and depressive mood, with higher total scores indicating greater emotional stress. For the present study, the primary outcome was the change in the total stress score from pre-to post-stimulation. Subscale scores were also explored to characterize specific aspects of stress reactivity.

In addition to stress assessment, we evaluated tDCS-related safety and tolerability using a brief questionnaire adapted from previous neuromodulation safety reports (Brunoni et al., 2011; Poreisz et al., 2007; Thair et al., 2017). Immediately after stimulation, participants rated the presence and intensity of common tDCS-related sensations, including itching, tingling, burning sensation, skin pain, headache, difficulties in concentrating, unpleasant sensations, nervousness, fatigue, drowsiness, changes in visual perception, and visual phenomena associated with the start or end of stimulation. For each sensation, participants reported whether it occurred during stimulation, after stimulation, or both. These safety data were examined descriptively to confirm that the stimulation protocol was well tolerated and to compare the incidence and severity of side effects between the tDCS and sham groups. Statistical analyses were performed in Python 3.

## 3. Results

### 3.1. fNIRS: Prefrontal Hemodynamic Changes and Connectivity

#### 3.1.1. Block Averaging

A two-way repeated-measures ANOVA with Group (tDCS, sham) as a between-subject factor and Block (prestim, durstim, poststim) as a within-subject factor was conducted for each channel. The Group × Block interaction was marginally significant for channel 1, *F*(2, 48) = 2.45, *p*_*uncorrected*_ =.097, and channel 5, *F*(2, 52) = 2.85, *p*_*uncorr*_ =.067, indicating a trend toward differential block-related changes between groups for these channels (Fig. 2). A significant main effect of Group was observed for channel 1, *F*(1, 48) = 4.27, *p*_*uncorr*_ =.50, with overall lower HbO levels in the tDCS group across blocks. A significant main effect of Block emerged for channels 8 (*F*(2, 44) = 3.55, *p*_*uncorr*_ =.037), 10 (*F*(2, 54) = 4.73, *p*_*uncorr*_ =.013), and 15 (*F*(2, 18) = 9.49, *p*_*FDR-corrected*_ =.023), suggesting consistent block-related modulation irrespective of group.

**Figure 2.**
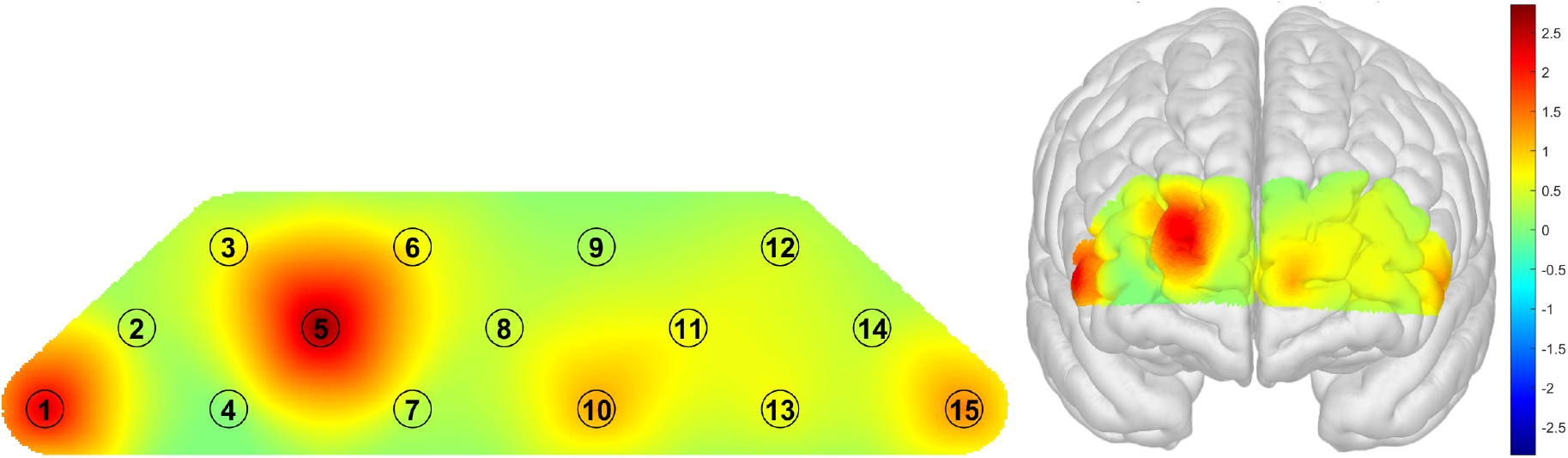
Results of two-way repeated measures ANOVA

Follow-up one-way repeated-measures ANOVAs within each group revealed that, in the sham group, only channel 1 showed a significant block effect, *F*(2, 16) = 4.74, *p*_*uncorr*_ =.024. In the tDCS group, significant block effects were found for channels 10 (*F*(2, 38) = 3.26, *p*_*uncorr*_ =.049), 11 (*F*(2, 40) = 4.23, *p*_*uncorr*_ =.022), and 15 (*F*(2, 10) = 14.23, *p*_*FDR-corr*_ =.018), see Fig. 3.

**Figure 3.**
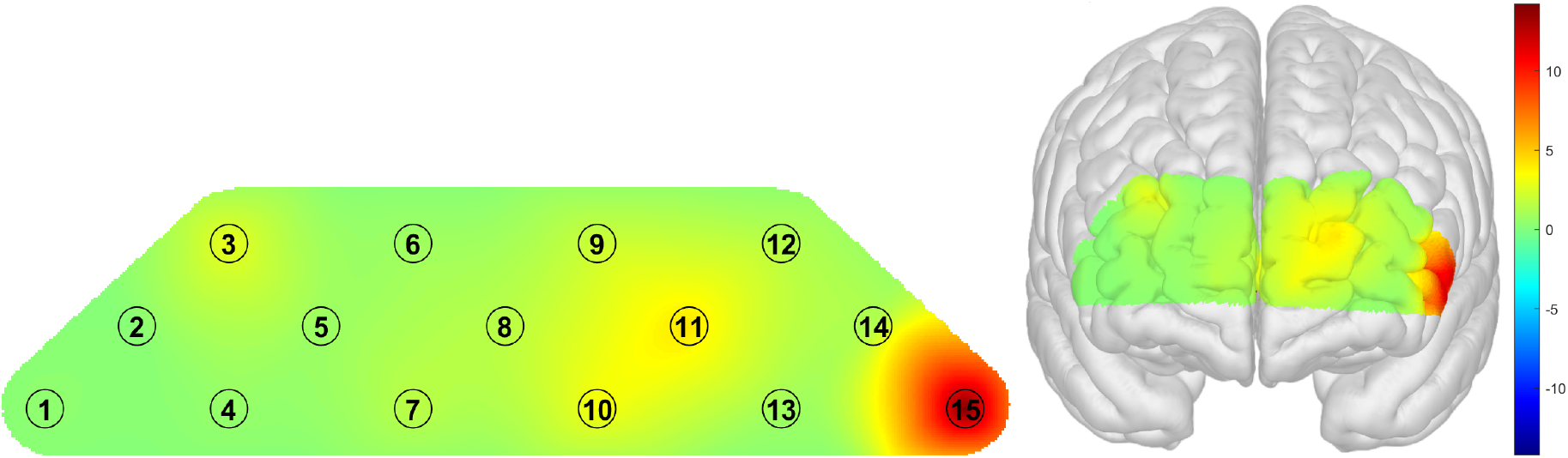
Results of one-way repeated measures ANOVA (tDCS group)

Planned pairwise *t*-tests comparing HbO levels between blocks (prestim vs. durstim, durstim vs. poststim, and poststim vs. prestim) were conducted separately for each group. In the sham group, significant differences were observed between prestim and durstim for channel 1 (*t*(7) = -2.97, *p*_*uncorr*_ =.021), and 12 (*t*(6) = -6.50, *p*_*FDR-corr*_ =.009), as well as between prestim and poststim for channel 12 (*t*(7) = -3.80, *p*_*uncorr*_ =.007). Given that participants in the sham condition did not receive any stimulation, these changes may reflect spontaneous fluctuations in prefrontal activation, potentially related to intrinsic activity within the default mode network.

In the tDCS group, HbO levels significantly decreased from prestim to durstim for channels 11 (*t*(*16*) = -2.34, *p*_*uncorr*_ =.033) and 15 (*t*(*5*) = -3.16, *p*_*uncorr*_ =.025). From durstim to poststim, significant decreases were observed in channels 5 (*t*(17) = -3.41, *p*_*FDR-corr*_ =.021), 6 (*t*(19) = -3.26, *p*_*FDR-corrected*_ =.021), 9 (*t*(20) = -2.76, *p*_*FDR-corr*_ =.045), and 12 (*t*(18) = -3.64, *p*_*FDR-corr*_ =.021).

Comparisons between prestim and poststim revealed significant differences for channels 2 (*t*(16) = -2.29, *p*_*uncorr*_ =.036), 3 (*t*(19) = -2.17, *p*_*uncorr*_ =.043), and 15 (*t*(5) = -6.29, *p*_*FDR-corr*_ =.022). Overall, the tDCS group exhibited stimulation-related decreases in prefrontal HbO during stimulation, followed by post-stimulation decreases toward or below baseline in specific regions, whereas changes in the sham group likely reflect spontaneous, non-stimulation-related neural activity.

Independent-samples *t*-tests on block difference scores (during minus pre, post minus during, and post minus pre) were conducted to compare the magnitude of change between the tDCS and sham groups for each channel. For the during minus pre contrast, a significant group difference was observed for channel 5 (*t*(25) = 2.52, *p* =.019), with the tDCS group showing larger HbO changes than the sham group. For the post minus during contrast, significant differences were found for channels 1 (*t*(27) = -2.23, *p* =.034), 6 (*t*(26) = -2..23, *p* =.035), and 12 (*t*(24) = -3.18, *p* <.01), with the tDCS group showing smaller HbO changes than the sham group, suggesting that the effects of tDCS persisted after stimulation. No significant group differences emerged for the post minus pre in any channel (all *ps* >.05).

#### 3.1.2. Connectivity

Connectivity analyses were conducted using a connectivity-based channel clustering approach. Channels were grouped into anatomically and functionally relevant clusters based on their spatial proximity and presumed cortical region coverage. This clustering allowed for the assessment of broader network-level changes in prefrontal connectivity, reducing the number of multiple comparisons and increasing interpretability of the results. Figure 4 illustrates the spatial layout of the channels and their assigned clusters.

**Figure 4.**
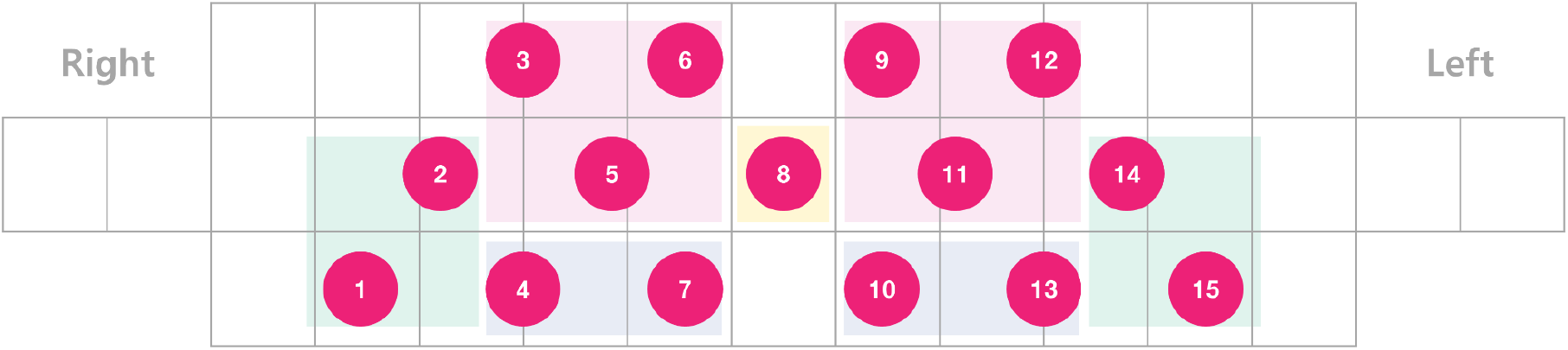
Channel configuration and connectivity-based clustering used for fNIRS connectivity analyses.

A two-way repeated-measures ANOVA with Group (tDCS, sham) as a between-subject factor and Block (prestim, durstim, poststim) as a within-subject factor was conducted for each clustering pair. No Group × Block interactions reached statistical significance after FDR correction, although marginal trends were observed for Upper Right - Center pair, *F*(2, 50) = 2.27, *p*_*uncorr*_ =.079. Significant main effects of Group were observed for 3 pairs (Lateral Left - Lateral Right, Lateral Right - Upper Left, and Lateral Right - Upper Right). Significant main effects of Block were found for 18 pairs (e.g., Lateral Left - Upper Left, Lateral Left - Upper Right, Lateral Left - Lower Left, Lateral Left - Lower Right, and etc.), indicating block-related modulation irrespective of group (Supplementary Table S1 for the full list and statistics).

Follow-up one-way repeated-measures ANOVAs conducted separately within each group revealed that: In the tDCS group, significant block effects were observed for 18 pairs, all *p*_*FDR-corr*_ <.05 (Table S2 for results). These effects were characterized by pronounced changes in interhemispheric connectivity, particularly for connections spanning homologous left–right regions, suggesting that tDCS may engage or reorganize bilateral network interactions. Significant block effects were observed across all major lateral regions, indicating potential modulation of coordination or synchronization between central and peripheral cortical areas. In contrast, the sham group showed a significant block effect only for the Upper Right - Lower Right pair, *F*(2, 18) = 8.78, *p*_*FDR-corr*_ =.046. The greater number and distribution of significant effects in the tDCS group, compared to sham, supports the interpretation that tDCS induced a robust modulation of neural connectivity beyond spontaneous fluctuations.

Planned pairwise *t*-tests on block difference scores (prestim vs. durstim, durstim vs. poststim, and prestim vs. poststim) revealed that, in the tDCS group, connectivity significantly decreased from prestim to durstim for seven pairs (all *p*_*uncorr*_ <.05), suggesting stimulation-induced decoupling between lateral and central regions. Connectivity also significantly decreased from durstim to poststim for four pairs (all *p*_*uncorr*_ <.05). Comparisons between prestim and poststim further indicated significant decreases for 14 pairs (*p*_*FDR-corr*_ <.05) (Table S3 for details). In contrast, the sham group showed a significant change from prestim to durstim only for the Lower Left–Center pair (*p*_*uncorr*_ <.05), from durstim to poststim for two pairs (*p*_*uncorr*_ <.05), and from prestim to poststim for two pairs (*p*_*FDR-corr*_ <.05).

Independent-samples *t*-tests on block difference scores revealed significant group differences. For during minus prestim contrast: Lateral Right - Center pair showed a significant effect, *t*(23) = -2.28, *p*_*uncorr*_ =.032, with the tDCS group exhibiting smaller connectivity changes than the sham group. For the post minus durstim contrast, the Upper Right - Lower Right pair was significant, *t*(28) = 2.52, *p*_*uncorr*_ =.032, indicating that the tDCS group demonstrated larger connectivity changes than the sham group. For the post minus prestim contrast, significant effects were observed for the Lateral Left - Lower Left pair, *t*(11) = -3.01, *p*_*uncorr*_ =.012, and the Upper Right - Center pair, *t*(24) = 2.33, *p*_*uncorr*_ =.028. These findings suggest that anodal stimulation over the left hemisphere inhibited connectivity among left lateral regions, whereas cathodal stimulation over the right hemisphere enhanced central network interactions after stimulation.

Taken together, the connectivity analyses indicate that low-intensity tDCS induced widespread and systematic modulation of prefrontal network dynamics, particularly evident in interhemispheric and lateral–central connections. While the sham group showed only limited and sporadic fluctuations, the tDCS group demonstrated consistent block-related changes across a broader set of connections, suggesting stimulation-specific decoupling and reorganization of functional networks. These results support the view that tDCS effects extend beyond focal cortical activation to influence large-scale network coordination, reflecting both inhibitory and facilitatory processes across bilateral prefrontal regions.

### 3.2. Heart Rate Variability (HRV)

We examined HRV metrics (RMSSD, SDNN, pNN50, LF, HF, LF/HF ratio, SD1, SD2, ApEn, DFA-α1) and mean heart rate across experimental blocks (prestim, durstim, poststim) in both the sham and tDCS groups.

A series of 2 (Group: sham, tDCS) × 3 (Block: prestim, durstim, poststim) repeated-measures ANOVAs were conducted for each metric. Significant main effects of Block were observed for parasympathetic-related indices including RMSSD (*F*(1.94, 31.06) = 6.27, *p* =.006), SDNN (*F*(1.99, 31.86) = 7.21, *p* =.003), pNN50 (*F*(1.95, 31.25) = 5.49, *p* =.009), as well as SD1 (*F*(1.97, 31.56) = 5.85, *p* =.007) and SD2 (*F*(1.99, 31.92) = 7.62, *p* =.002) (Fig. 5). These effects indicate that autonomic regulation varied significantly across stimulation phases. In contrast, no significant Group × Block interactions were detected for any metric, suggesting that both groups followed broadly similar block-related patterns. However, heart rate showed a marginally significant main effect of Group, *F*(1, 16) = 3.38, *p* =.085, with a trend toward higher mean heart rates in the sham group compared to the tDCS group across blocks. Follow-up contrasts indicated that sham participants had higher heart rates than the tDCS group for the poststim block, *t*(16) = 2.34, *p* =.033, whereas no significant differences emerged during prestim or durstim.

**Figure 5.**
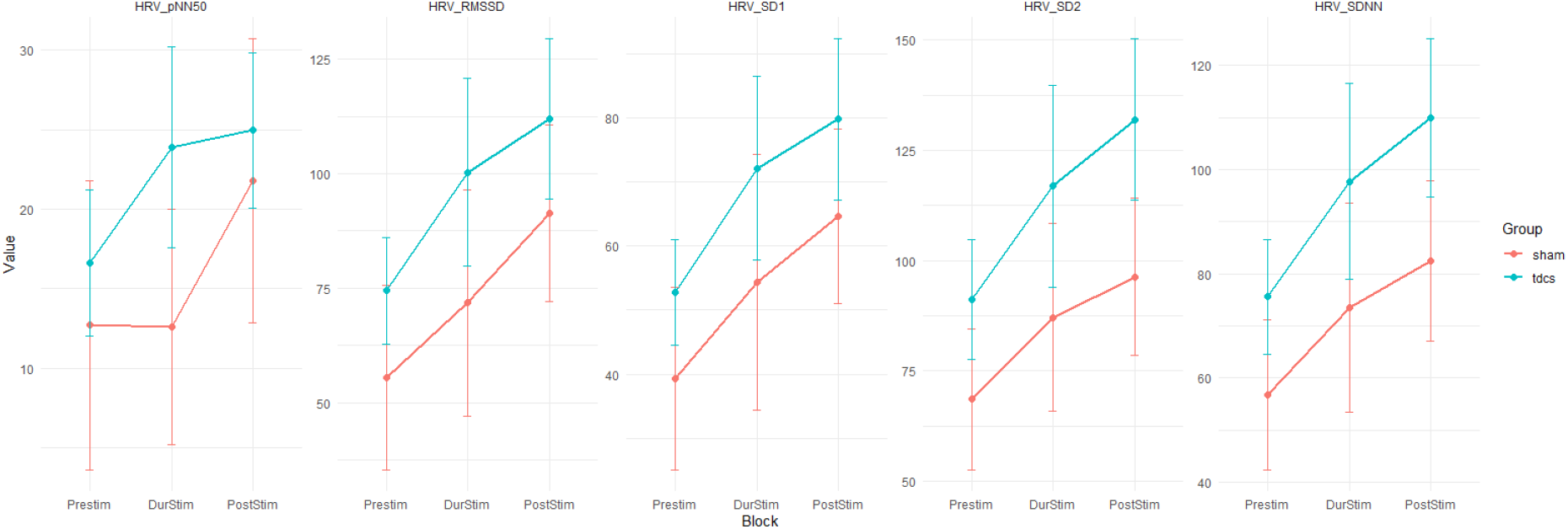
Group-level changes in parasympathetic HRV measures (RMSSD, SDNN, pNN50, SD1, SD2) across experimental blocks

Follow-up one-way repeated-measures ANOVAs within each group revealed that the tDCS group exhibited significant block effects for RMSSD (*F*(2, 22) = 4.60, *p* =.021), SDNN (*F*(2, 22) = 6.64, *p* =.006), pNN50 (*F*(2, 22) = 4.40, *p* =.025), SD1 (*F*(2, 22) = 4.73, *p* =.020), and SD2 (*F*(2, 22) = 7.11, *p* =.004) (Fig. 6). Planned pairwise comparisons (Bonferroni-corrected) indicated that these changes were primarily driven by increases from prestim to poststim. For example, SDNN significantly increased, *t*(11) = -3.34, *p* =.02, and SD2 also showed a significant increase, *t*(11) = -3.52, *p* =.014, *d* = 0.91. Similar patterns were observed for RMSSD (*p* =.06, trend-level), pNN50 (*t*(11) = -3.03, *p* =.034), and SD1 (*p* =.066, trend-level). These findings are consistent with stimulation-related increases in parasympathetic tone, as indexed by time-domain HRV measures (Brunoni et al., 2013; Ko et al., 2024). In contrast, the sham group showed no reliable block effects after correction, although marginal changes were observed for RMSSD and pNN50. No significant block effects emerged for frequency-domain indices (LF, HF, LF/HF), ApEn, DFA-α_1_, or mean Heart Rate in either group.

**Figure 6.**
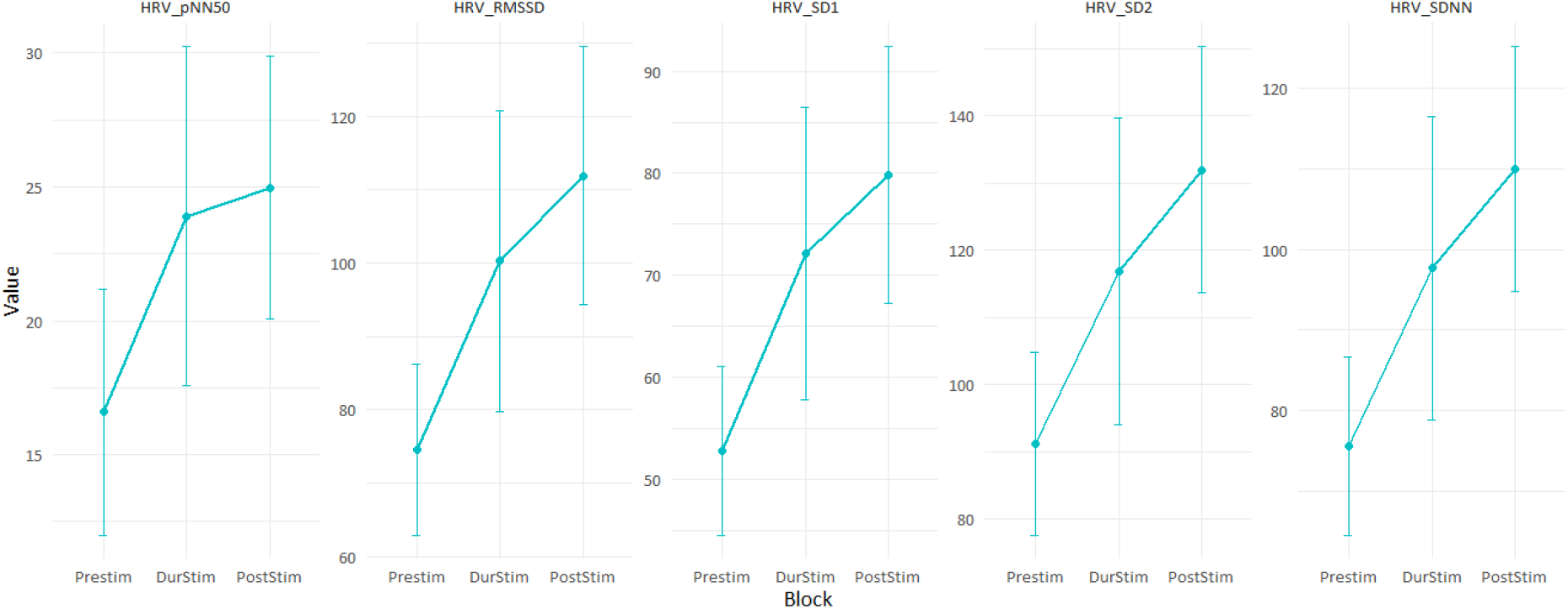
Block-wise changes in parasympathetic HRV measures in the tDCS group

Independent-samples t-tests comparing sham and tDCS groups at each block revealed no significant between-group differences for any HRV index (all *ps* >.05), though heart rate tended to be marginally lower in the tDCS group during poststim, t(7.46) = 2.06, p =.076.

Taken together, these results suggest that tDCS was associated with modest increases in parasympathetic activity (RMSSD, SDNN, pNN50, SD1, SD2) from prestim to poststim, whereas the sham group did not show systematic changes beyond natural fluctuations. Although no robust Group × Block interactions emerged, within-group patterns indicate that tDCS may exert subtle modulatory effects on autonomic regulation.

### 3.3. Phase Locking Value (PLV)

We examined phase synchronization between prefrontal HbO and the slow-band PPG envelope. A mixed ANOVA with Group (tDCS, sham) as a between-subject factor and Block (prestim, durstim, poststim) as a within-subject factor on PLV revealed no significant main effects or interactions: Group, *F*(1, 16) = 0.33, *p* =.573, partial η^2^ =.020; Block, *F*(2, 32) = 1.38, *p* =.266, partial η^2^ =.079 (Greenhouse–Geisser ε ≈ 0.857); Group × Block, *F*(2, 32) = 0.45, *p* =.641, partial η^2^ =.027. Consistent with the analysis, no channel × block between-group differences survived FDR correction. Group-wise descriptive trends are visualized in Fig. 7 (mean ± 95% CI per block).

**Figure 7.**
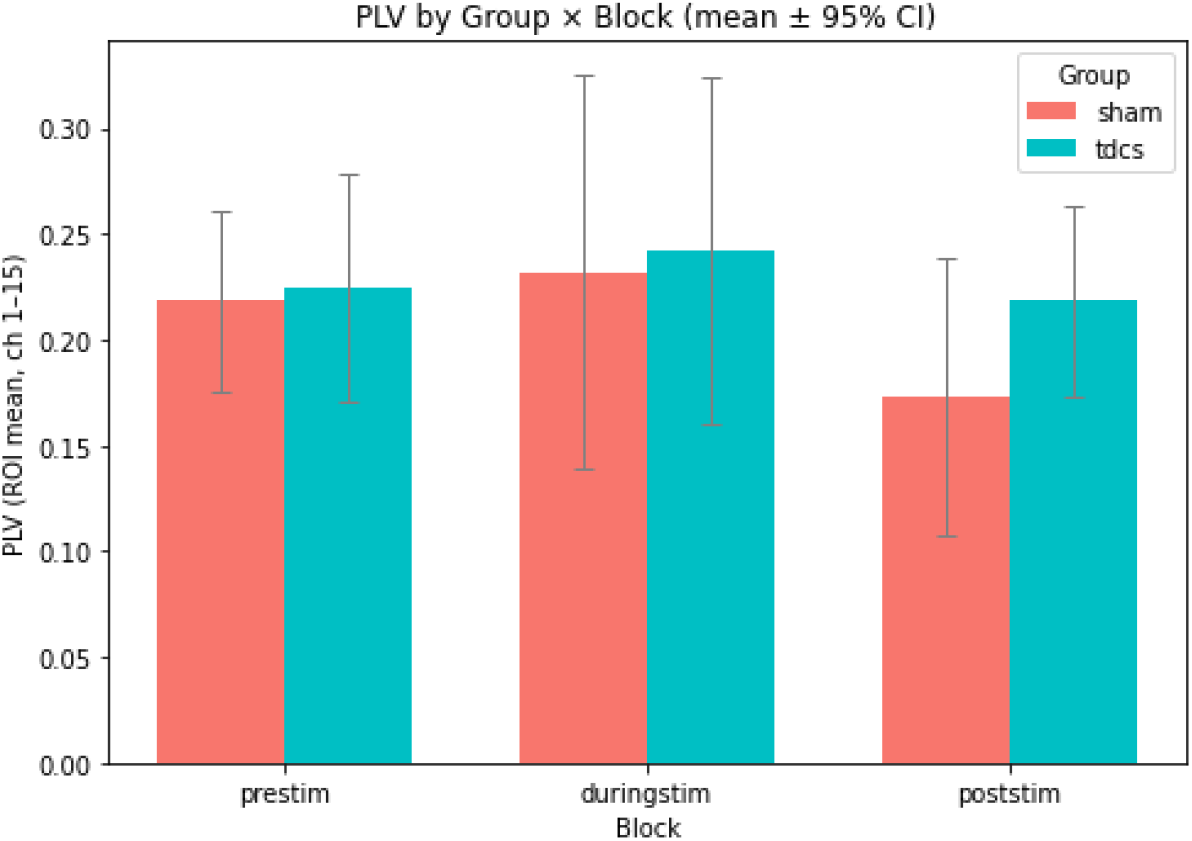
Group-wise PLV (Channels 1–15) across blocks; bars indicate mean ± 95% CI.

One-way repeated measures ANOVAs were conducted for each group. Within the sham group, the main effect of Block was not significant, F(2, 10) = 2.11, p =.172 (Greenhouse–Geisser: p-GG =.185). Similarly, the tDCS group showed no Block effect, F(2, 22) = 0.38, p =.685 (p-GG =.649). Continuing, we conducted blockwise between-group comparisons at the ROI level (mean of channels 1–15). During prestim, Levene’s test indicated unequal variances (p =.031), so we used Welch’s t; the difference was not significant, t = −0.21, p =.836, Hedges’ g = 0.08 (sham: M = 0.219; tDCS: M = 0.225; n = 6 vs. 12). In the durstim block, the equal-variance assumption held (Levene p =.177) and the independent-samples test showed no difference, t(16) = −0.18, p =.861, Hedges’ g = 0.09 (sham: M = 0.232; tDCS: M = 0.242). In the poststim block, variances were also homogeneous (Levene p =.883); the group difference was not significant, t(16) = −1.34, p =.199, Hedges’ g = 0.64 (sham: M = 0.173; tDCS: M = 0.219). After multiplicity control across blocks, all effects remained non-significant (Holm-adjusted p ≥.598; FDR-adjusted p ≥.598).

Overall, under the corrected slow-band PLV pipeline (time-base alignment to fNIRS; band-limited phase extraction), we did not observe reliable changes in cardio-cerebral phase coupling attributable to low-intensity tDCS. These results suggest that, at 0.07–0.15 Hz and with the present protocol, cardio-cerebral phase synchrony remained broadly comparable across groups and blocks.

### 3.4. Subjective Survey

Participants (*N* = 22; tDCS = 14, sham = 8) completed an emotional stress inventory (ESI) before and after the stimulation/sham session. This self-report scale included 6 ranges: state anger, state anxiety, state depression, trait anger, trait anxiety, and trait depression. At prestim, the tDCS group scored numerically higher than the sham group across most measures; however, only state depression differed significantly at baseline. All other pre-session differences were nonsignificant (all *p*s >.05) (Table 1).

**Table 1.** Pre-stimulation (baseline) group differences on self-report stress measures.

| Measure | $M_{\text{tDCS}}$ | $SD_{\text{tDCS}}$ | $M_{\text{sham}}$ | $SD_{\text{sham}}$ | $t$ | df | $p_{\text{FDR-corr}}$ | $d$ |
| --- | --- | --- | --- | --- | --- | --- | --- | --- |
| state anger | 10.64 | 4.50 | 8.50 | 3.12 | 1.31 | 10.9 | .21 | 0.53 |
| state anxiety | 18.93 | 7.73 | 14.25 | 8.05 | 1.33 | 15.2 | .21 | 0.60 |
| state depression | 19.57 | 6.63 | 12.13 | 4.55 | 3.11 | 10.9 | .03* | 1.24 |
| trait anger | 14.00 | 5.59 | 10.38 | 5.37 | 1.50 | 14.2 | .21 | 0.66 |
| trait anxiety | 23.64 | 7.00 | 19.38 | 7.43 | 1.32 | 15.5 | .21 | 0.60 |
| trait depression | 14.07 | 6.12 | 10.88 | 4.76 | 1.36 | 11.9 | .21 | 0.56 |

We tested whether self-reported symptoms changed differentially by group by comparing change scores (Post − Pre) between the tDCS and sham groups using Welch’s independent t tests (with two time points, this is equivalent to the Time × Group interaction). We then conducted follow-up ANCOVAs on post-session scores with the corresponding pre-session score entered as a covariate to account for baseline differences.

A significant Group × Time effect emerged for state depression (S-DEP): the tDCS group decreased more than sham (tDCS: *M* change = −3.43; sham: −0.38), *t*(20) = −2.21, *p* =.040, *d* = −0.90; however, this effect was no longer significant when pre-session S-DEP was included as a covariate (ANCOVA group *p* =.307), consistent with the sizable baseline imbalance on this scale. Trait anger (T-ANG) also showed a significant group difference in change (tDCS: −2.43; sham:

+1.75), *t*(20) = −2.93, *p* =.013 *d* = −1.40. This effect remained after controlling for baseline in an ANCOVA on post scores (*p* <.01). State anger displayed a trend toward a larger reduction in the tDCS group (tDCS: −1.86; sham: +1.75), *t*(20) = −2.12, *p* =.057, *d* = −1.02; the corresponding ANCOVA showed a similar trend (*p* =.080). For State anxiety (S-ANX), Trait depression (T-DEP), and trait anxiety (T-ANX), group differences in change were not significant (all *p*s ≥.121), and ANCOVAs on post scores controlling for baseline were likewise nonsignificant (S-ANX *p* =.149; T-DEP *p* =.431; T-ANX *p* =.821). Collectively, the results indicate selective improvements following tDCS—most robust for anger-related symptoms—while other domains showed no reliable group-specific change beyond baseline levels (Fig. 8).

**Figure 8.**
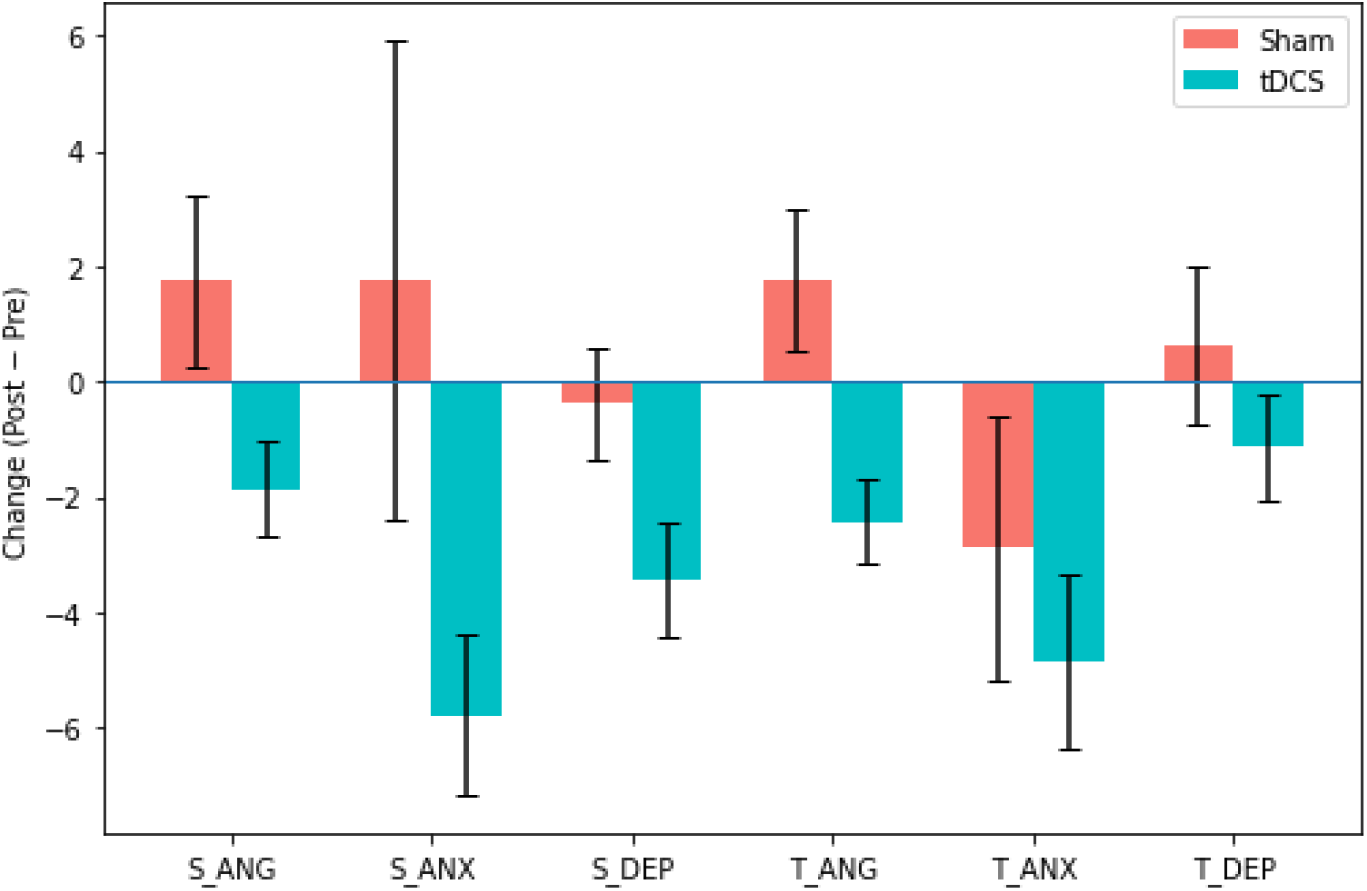
Group differences in change scores (Post − Pre) across all measures.

Paired *t* tests showed that the tDCS group reported significant reductions on multiple scales (df = 13): S-ANX, *t* = −4.18, *p* =.001, *dz* = −1.12; S-ANG, *t* = −2.22, *p* =.045, *dz* = −0.59; T-ANG, *t* = −3.39, *p* =.005, *dz* = −0.91; T-ANX, *t* = −3.19, *p* =.007, *dz* = −0.85; S-DEP, *t* = −3.45, *p* =.004, *dz* = −0.92. In contrast, the sham group showed no significant pre–post changes on any scale (df = 7; all *p*s ≥.20; e.g., S-ANX, *t* = 0.42, *p* =.686, *dz* = 0.15; T-ANG, *t* = 1.42, *p* =.200, *dz* = 0.50).

Together, these findings suggest selective self-reported improvements following tDCS, most consistently for anger-related scales—most notably T-ANG, which remained significant after adjustment for baseline—and to a lesser extent for S-DEP (an effect attenuated when baseline differences were controlled). Other domains (S-ANX, T-DEP, T-ANX) showed no reliable Group × Time effects. Given the modest sample (*n*_sham_ = 8, *n*_tDCS_ = 14) and a notable baseline difference on S-DEP, the results should be interpreted cautiously; nevertheless, they accord with prior work indicating that a single-session prefrontal neuromodulation in healthy individuals yields little to no immediate change in mood (Morgan et al., 2014; Remue et al., 2016). Thus, any physiological effects associated with the present stimulation were likely too subtle to produce overt changes in self-perceived state over this short time frame.

Immediately after the stimulation session, participants completed a 12-item adverse-effects checklist that asked them to rate sensations experienced during stimulation and after the session. A composite severity score (mean across items) showed no between-group differences at either time point (During: *t*(15.6) = −0.39, *p* =.699, *d* = −0.16; Post: *t*(21.1) = −0.01, *p* =.996, *d* ≈ 0; *n*_*sham*_ = 10, *n*_*tDCS*_ = 22). Within groups, ratings decreased from during to post, indicating that sensations were transient (Sham: *t*(9) = −4.04, *p* =.003, *dz* = 1.28; tDCS: *t*(21) = −4.02, *p* <.001, *dz* = 0.86).

Item-level tests likewise revealed no between-group differences after FDR correction at either time point, while several common sensations (e.g., tingling, skin pain, and changes in visual perception) declined from during to post within groups (FDR-corrected *p*s <.05), with no items increasing. Together, these results indicate that the stimulation protocol was well tolerated, producing only mild, short-lived sensations and no evidence of excess adverse effects relative to sham.

## 4. Discussion

This study provides new evidence that low-intensity bifrontal tDCS can simultaneously influence cortical, autonomic, and affective domains in healthy humans. We found that anodal stimulation over the left DLPFC (with a 0.375 mA current) led to decreased oxygenated hemoglobin (HbO) in the underlying cortex during stimulation, altered prefrontal functional connectivity patterns, increased vagally mediated heart rate variability, and a post-stimulation reduction in self-reported anger, compared to a sham condition. Taken together, these findings support the notion that even sub-threshold levels of direct current can engage neurovascular and neurovisceral mechanisms, producing a coordinated shift toward a calmer physiological and emotional state. In the following, we discuss each of these outcomes in the context of known neurobiology and prior research, highlighting potential mechanisms of cortical-autonomic coupling and the implications for neuromodulation science.

The reduction in oxyhemoglobin under the anode was unexpected, as anodal stimulation is usually linked to increased excitability and metabolic demand. However, paradoxical decreases have been reported in other tDCS-neuroimaging studies, particularly at rest (Figeys et al., 2021; Stagg et al., 2013). One explanation is that weak stimulation preferentially recruits inhibitory interneurons or triggers homeostatic downregulation of spontaneous activity, resulting in lower blood flow (Stagg & Nitsche, 2011). Another possibility is that direct current affected vascular tone, leading to perfusion changes independent of neural firing (Campos et al., 2024). These interpretations highlight that hemodynamic responses to tDCS are state- and context-dependent, and that decreased HbO should not be automatically viewed as negative.

At the network level, tDCS altered functional connectivity patterns within the prefrontal cortex. We observed reduced synchronization across left frontal sites and reorganization of bilateral connections, consistent with prior fNIRS and fMRI findings (Bouchard et al., 2023; Keeser et al., 2011). Rather than uniformly increasing connectivity, tDCS appears to rebalance interactions, strengthening some connections while weakening others. Such reconfiguration suggests that weak current may act less as a focal activator and more as a modulator of network dynamics, nudging large-scale circuits into a different equilibrium. These network changes are theoretically important because prefrontal connectivity underlies executive control and emotion regulation. Reduced intra-frontal coupling might reflect more efficient, segregated processing, while increased interhemispheric or lateral–central connectivity could support integration of regulatory signals (Bouchard et al., 2023). Although we did not include a cognitive task, such network modulation could provide the substrate for improved executive or affective outcomes seen in other tDCS studies.

Autonomic outcomes strongly complemented the neural effects. Time-domain HRV measures (RMSSD, SDNN, pNN50, SD1, SD2) increased significantly with stimulation, indicating enhanced parasympathetic influence. Heart rate tended to decline, further supporting a vagal shift. These results align with the neurovisceral integration model, in which prefrontal activity exerts inhibitory control over subcortical autonomic centers to maintain flexible cardiovascular regulation (Thayer & Lane, 2009; Nikolin et al., 2017). By increasing DLPFC excitability, tDCS likely amplified this “vagal brake,” producing a measurable change in heart rhythm. That low-intensity stimulation produced clear autonomic effects is notable. Prior work has demonstrated HRV modulation with higher currents (Gu et al., 2022; Ko et al., 2024), but our results suggest that even sub-milliamp currents can influence central autonomic pathways. This raises the possibility that cortical-autonomic circuits are highly sensitive to subtle perturbations, and that clinical effects might be achievable with lower, safer, and more tolerable doses of stimulation.

Emotional stress data provided converging evidence. Active tDCS led to reductions in state and trait anger, while other mood domains remained unchanged. Although modest and in need of replication, this selective effect suggests that tDCS induced a calming influence most evident in anger regulation. One likely mechanism is that increased vagal tone translated into subjective feelings of reduced irritability (Shaffer & Ginsberg, 2017). Alternatively, heightened prefrontal control may have directly suppressed limbic circuits associated with anger (Carnevali et al., 2020). Either way, the convergence of neural, autonomic, and affective outcomes points to an integrated regulatory shift.

Interestingly, reductions were observed not only in state anger but also in trait anger, usually considered a stable disposition. This likely reflects a transient shift in self-perception rather than a durable personality change. Still, the finding underscores that even a single low-dose session can influence how individuals appraise their emotional tendencies. Context may have played a role: participants were resting quietly, which may have allowed subtle effects of stimulation to emerge without interference from task demands (Morgan et al., 2014).

The simultaneous occurrence of decreased DLPFC HbO, increased HRV, and reduced anger suggests cortical-autonomic coupling. Modulating the DLPFC appears to have triggered descending pathways that coordinated cardiovascular and emotional regulation. From an allostatic perspective, the brain responded to the perturbation of tDCS by adjusting body state toward parasympathetic dominance, consistent with a relaxation response (Thayer et al., 2012; Silvani et al., 2016). This systemic rebalancing highlights the brain–heart axis as a key mediator of neuromodulation effects.

Our findings also carry methodological implications. They demonstrate the value of multimodal measurement: only by combining fNIRS, HRV, and self-reports could we capture the full cascade of changes. Had we relied on a single measure, the picture would have been incomplete. This supports calls for integrated approaches in neuromodulation research to better characterize brain–body interactions. Clinically, the ability of tDCS to increase HRV and reduce anger points toward potential applications in stress-related and affective disorders. Low HRV is a risk factor for poor emotion regulation and cardiovascular disease, and excessive anger is a feature of many psychiatric conditions. Enhancing prefrontal–autonomic coupling with non-invasive, low-intensity stimulation could offer a safe adjunct to therapy. Moreover, the demonstration that ultra-low currents are effective suggests that tolerability can be improved without sacrificing efficacy (Gomez-Alvaro et al., 2024).

At the same time, the study has limitations. The sample was small and restricted to young men, limiting generalizability. Effects were assessed only acutely, so it remains unclear how long they persist or whether repeated sessions would produce cumulative benefits. fNIRS measured only superficial cortical signals, and autonomic measures were limited to HRV without direct assessment of sympathetic markers such as blood pressure or skin conductance. Mood findings, particularly on trait anger, should be interpreted cautiously given the modest sample size and potential baseline differences.

Future research should replicate these findings in larger, more diverse samples, explore dose - response relationships across current intensities, and examine durability of effects across repeated sessions. Adding complementary measures such as EEG, fMRI, or respiratory monitoring would help disentangle neural from systemic contributions. Clinical studies could test whether low-intensity tDCS reduces stress reactivity, improves mood regulation, or benefits patients with autonomic dysregulation.

In summary, this study demonstrates that low-intensity bifrontal tDCS can engage prefrontal neurovascular and autonomic systems in tandem, producing physiological and emotional signatures of relaxation. The results reinforce the idea that the brain and body respond as an integrated unit to neuromodulation, and that even weak stimulation can tip this system toward balance. By illuminating cortical-autonomic coupling, our findings contribute to a systemic view of tDCS and its potential for enhancing emotional and physiological regulation.

## 5. Conclusions

In conclusion, this study shows that even ultra-low intensity bifrontal tDCS (0.375 mA) can modulate brain and body function at rest. Anodal stimulation over the left DLPFC reduced local oxyhemoglobin, altered prefrontal connectivity, enhanced vagally mediated heart rate variability, and selectively reduced anger in healthy men. These results support the view that weak direct current engages integrated neurovascular and neurovisceral mechanisms, producing a coordinated shift toward a calmer physiological and emotional state.

Beyond these immediate effects, our findings highlight the DLPFC’s role as a cortical hub linking neural activity to autonomic regulation. Multimodal monitoring revealed that cortical changes were mirrored by autonomic and affective outcomes, underscoring the value of assessing brain–body interactions in neuromodulation research. For clinical translation, the results suggest that low-intensity stimulation could offer a safe, well-tolerated strategy to promote autonomic balance and emotional regulation, with potential applications in stress-related and anger-prone conditions.

Overall, this work illustrates the systemic reach of neuromodulation: a small current applied to the scalp can reshape cortical activity, cardiac dynamics, and mood. Future research should determine optimal parameters, examine persistence across repeated sessions, and test efficacy in clinical populations. By advancing our understanding of cortical-autonomic coupling, low-intensity tDCS may emerge as a versatile tool for neuroscience and a promising adjunct in interventions targeting both mental and physical health.

## Acknowledgments

We sincerely thank the participants for their time and commitment to this study, as well as our research staff for their assistance with data collection and management.

