## Supplementary material for "Transcranial Direct Current Stimulation Modulates Resting Brain Hemodynamics and Autonomic Function: A Multimodal fNIRS-HRV Study": Table S1

**Appendix**

**Table S1**. Two-way Repeated-Measures ANOVA Results for Connectivity-Based Channel Clusters (HbO)

| ROI | Effects | *F* | df1 | df2 | $p_{uncorr}$ | $p_{FDR\text{-}corr}$ |
|---|---|---|---|---|---|---|
| Lateral_Left - Lateral_Right | Group | 5.40 | 1 | 22 | 0.04* | 0.26 |
| | Block | 3.13 | 2 | 22 | 0.06 | 0.07 |
| | Group*Block | 1.92 | 2 | 22 | 0.17 | 0.61 |
| Lateral_Left - Upper_Left | Group | 1.28 | 1 | 22 | 0.28 | 0.40 |
| | Block | 5.20 | 2 | 22 | 0.01* | 0.02* |
| | Group*Block | 0.39 | 2 | 22 | 0.68 | 0.85 |
| Lateral_Left - Upper_Right | Group | 2.01 | 1 | 22 | 0.18 | 0.32 |
| | Block | 4.75 | 2 | 22 | 0.02* | 0.03* |
| | Group*Block | 0.46 | 2 | 22 | 0.64 | 0.85 |
| Lateral_Left - Lower_Left | Group | 2.73 | 1 | 22 | 0.13 | 0.30 |
| | Block | 3.90 | 2 | 22 | 0.04* | 0.04* |
| | Group*Block | 1.97 | 2 | 22 | 0.16 | 0.61 |
| Lateral_Left - Lower_Right | Group | 3.09 | 1 | 22 | 0.11 | 0.30 |
| | Block | 5.35 | 2 | 22 | 0.01* | 0.02* |
| | Group*Block | 2.56 | 2 | 22 | 0.10 | 0.61 |
| Lateral_Left - Center | Group | 1.46 | 1 | 18 | 0.26 | 0.40 |
| | Block | 3.01 | 2 | 18 | 0.07 | 0.08 |
| | Group*Block | 1.35 | 2 | 18 | 0.28 | 0.66 |
| Lateral_Right - Upper_Left | Group | 8.58 | 1 | 58 | 0.01** | 0.14 |
| | Block | 3.93 | 2 | 58 | 0.03* | 0.03* |
| | Group*Block | 0.73 | 2 | 58 | 0.49 | 0.85 |
| Lateral_Right - Upper_Right | Group | 4.21 | 1 | 58 | 0.05* | 0.26 |
| | Block | 5.97 | 2 | 58 | 0.00** | 0.01** |
| | Group*Block | 0.83 | 2 | 58 | 0.44 | 0.85 |
| Lateral_Right - Lower_Left | Group | 3.83 | 1 | 54 | 0.06 | 0.26 |
| | Block | 5.97 | 2 | 54 | 0.00** | 0.01** |
| | Group*Block | 1.57 | 2 | 54 | 0.22 | 0.65 |
| Lateral_Right - Lower_Right | Group | 2.65 | 1 | 56 | 0.11 | 0.30 |
| | Block | 5.18 | 2 | 56 | 0.01** | 0.02* |
| | Group*Block | 0.45 | 2 | 56 | 0.64 | 0.85 |
| Lateral_Right - Center | Group | 1.19 | 1 | 48 | 0.29 | 0.40 |
| | Block | 2.69 | 2 | 48 | 0.08 | 0.08 |
| | Group*Block | 2.02 | 2 | 48 | 0.14 | 0.61 |
| Upper_Left - Upper_Right | Group | 4.00 | 1 | 60 | 0.05 | 0.26 |
| | Block | 7.09 | 2 | 60 | 0.00** | 0.01** |
| | Group*Block | 0.44 | 2 | 60 | 0.65 | 0.85 |
| Upper_Left - Lower_Left | Group | 1.94 | 1 | 56 | 0.17 | 0.32 |
| | Block | 6.20 | 2 | 56 | 0.00** | 0.01** |
| | Group*Block | 0.19 | 2 | 56 | 0.83 | 0.85 |
| Upper_Left - Lower_Right | Group | 1.99 | 1 | 58 | 0.17 | 0.32 |
| | Block | 5.30 | 2 | 58 | 0.01** | 0.01* |
| | Group*Block | 0.17 | 2 | 58 | 0.85 | 0.85 |
| Upper_Left - Center | Group | 0.55 | 1 | 50 | 0.47 | 0.58 |
| | Block | 10.67 | 2 | 50 | 0.00*** | 0.00** |
| | Group*Block | 1.33 | 2 | 50 | 0.27 | 0.66 |
| Upper_Right - Lower_Left | Group | 0.01 | 1 | 56 | 0.93 | 0.93 |
| | Block | 3.94 | 2 | 56 | 0.02* | 0.03* |

| | | | | | | |
|---|---|---|---|---|---|---|
| | Group*Block | 0.19 | 2 | 56 | 0.82 | 0.85 |
| Upper_Right - Lower_Right | Group | 0.21 | 1 | 58 | 0.65 | 0.72 |
| | Block | 8.63 | 2 | 58 | 0.00*** | 0.01** |
| | Group*Block | 1.80 | 2 | 58 | 0.17 | 0.61 |
| Upper_Right - Center | Group | 0.01 | 1 | 50 | 0.92 | 0.93 |
| | Block | 7.87 | 2 | 50 | 0.00** | 0.01** |
| | Group*Block | 2.67 | 2 | 50 | 0.08 | 0.61 |
| Lower_Left - Lower_Right | Group | 2.45 | 1 | 54 | 0.13 | 0.30 |
| | Block | 7.71 | 2 | 54 | 0.00** | 0.01** |
| | Group*Block | 0.38 | 2 | 54 | 0.69 | 0.85 |
| Lower_Left - Center | Group | 0.83 | 1 | 46 | 0.37 | 0.49 |
| | Block | 6.28 | 2 | 46 | 0.00** | 0.01** |
| | Group*Block | 0.59 | 2 | 46 | 0.56 | 0.85 |
| Lower_Right - Center | Group | 0.28 | 1 | 48 | 0.60 | 0.70 |
| | Block | 6.55 | 2 | 48 | 0.00** | 0.01** |
| | Group*Block | 0.23 | 2 | 48 | 0.80 | 0.85 |

*Note*. df = degrees of freedom. * $p < .05$. ** $p < .01$. *** $p < .001$
