## Supplementary material for "Transcranial Direct Current Stimulation Modulates Resting Brain Hemodynamics and Autonomic Function: A Multimodal fNIRS-HRV Study": Table S2

**Table S2.** Results of follow-up one-way repeated-measures ANOVAs on connectivity-based channel clusters in the tDCS group.

| ROI | *F* | df1 | df2 | $p_{uncorr}$ | $p_{FDR\text{-}corr}$ |
|---|---|---|---|---|---|
| Lateral_Left - Lateral_Right | 6.04 | 2 | 12 | 0.02* | 0.03* |
| Lateral_Left - Upper_Left | 4.28 | 2 | 12 | 0.04* | 0.05* |
| Lateral_Left - Upper_Right | 3.97 | 2 | 8 | 0.06 | 0.07 |
| Lateral_Left - Lower_Left | 9.37 | 2 | 8 | 0.01** | 0.02* |
| Lateral_Left - Lower_Right | 10.02 | 2 | 12 | 0.00** | 0.01* |
| Lateral_Left - Center | 3.95 | 2 | 8 | 0.06 | 0.07 |
| Lateral_Right - Upper_Left | 4.75 | 2 | 38 | 0.01* | 0.03* |
| Lateral_Right - Upper_Right | 7.70 | 2 | 34 | 0.00** | 0.01* |
| Lateral_Right - Lower_Left | 12.56 | 2 | 30 | 0.00*** | 0.00** |
| Lateral_Right - Lower_Right | 4.55 | 2 | 38 | 0.02* | 0.03* |
| Lateral_Right - Center | 6.72 | 2 | 32 | 0.00** | 0.01* |
| Upper_Left - Upper_Right | 5.24 | 2 | 42 | 0.01** | 0.02* |
| Upper_Left - Lower_Left | 3.48 | 2 | 40 | 0.04* | 0.05* |
| Upper_Left - Lower_Right | 2.85 | 2 | 40 | 0.07 | 0.07 |
| Upper_Left - Center | 8.21 | 2 | 34 | 0.00** | 0.01* |
| Upper_Right - Lower_Left | 3.74 | 2 | 34 | 0.03* | 0.04* |
| Upper_Right - Lower_Right | 3.91 | 2 | 38 | 0.03* | 0.04* |
| Upper_Right - Center | 6.00 | 2 | 32 | 0.01** | 0.02* |
| Lower_Left - Lower_Right | 5.52 | 2 | 38 | 0.01** | 0.02* |
| Lower_Left - Center | 3.91 | 2 | 34 | 0.03* | 0.04* |
| Lower_Right - Center | 7.48 | 2 | 30 | 0.00** | 0.01* |

*Note*. df = degrees of freedom. * $p < .05$. ** $p < .01$. *** $p < .001$
