## Supplementary material for "Transcranial Direct Current Stimulation Modulates Resting Brain Hemodynamics and Autonomic Function: A Multimodal fNIRS-HRV Study": Table S3

**Table S3.** Planned pairwise *t*-tests on block difference scores (prestim vs. durstim, durstim vs. poststim, and prestim vs. poststim)

| Block Contrast | ROI | *t* | df | $p_{uncorr}$ | $p_{FDR\text{-}corr}$ |
|---|---|---|---|---|---|
| durstim - prestim | Lateral_Left - Lateral_Right | -3.28 | 6 | 0.02* | 0.07 |
| | Lateral_Left - Upper_Left | -2.09 | 6 | 0.08 | 0.16 |
| | Lateral_Left - Upper_Right | -2.37 | 6 | 0.06 | 0.12 |
| | Lateral_Left - Lower_Left | -1.97 | 6 | 0.10 | 0.17 |
| | Lateral_Left - Lower_Right | -4.99 | 6 | 0.00** | 0.05 |
| | Lateral_Left - Center | -3.58 | 4 | 0.02* | 0.08 |
| | Lateral_Right - Upper_Left | -1.50 | 20 | 0.15 | 0.22 |
| | Lateral_Right - Upper_Right | -2.06 | 20 | 0.05 | 0.12 |
| | Lateral_Right - Lower_Left | -2.75 | 19 | 0.01* | 0.07 |
| | Lateral_Right - Lower_Right | -1.23 | 18 | 0.24 | 0.31 |
| | Lateral_Right - Center | -2.75 | 16 | 0.01* | 0.07 |
| | Upper_Left - Upper_Right | -0.96 | 20 | 0.35 | 0.37 |
| | Upper_Left - Lower_Left | -1.13 | 20 | 0.27 | 0.34 |
| | Upper_Left - Lower_Right | -0.73 | 20 | 0.48 | 0.48 |
| | Upper_Left - Center | -2.30 | 18 | 0.03* | 0.10 |
| | Upper_Right - Lower_Left | -0.96 | 20 | 0.35 | 0.37 |
| | Upper_Right - Lower_Right | -1.06 | 18 | 0.30 | 0.35 |
| | Upper_Right - Center | -1.34 | 18 | 0.20 | 0.28 |
| | Lower_Left - Lower_Right | -1.62 | 19 | 0.12 | 0.20 |
| | Lower_Left - Center | -2.10 | 17 | 0.05 | 0.12 |
| | Lower_Right - Center | -2.72 | 17 | 0.01* | 0.07 |
| poststim - durstim | Lateral_Left - Lateral_Right | 0.00 | 6 | 1.00 | 1.00 |
| | Lateral_Left - Upper_Left | -1.55 | 6 | 0.17 | 0.40 |
| | Lateral_Left - Upper_Right | -1.14 | 6 | 0.30 | 0.45 |
| | Lateral_Left - Lower_Left | -3.50 | 5 | 0.02* | 0.12 |
| | Lateral_Left - Lower_Right | -0.47 | 6 | 0.66 | 0.77 |
| | Lateral_Left - Center | -0.20 | 4 | 0.85 | 0.94 |
| | Lateral_Right - Upper_Left | -2.66 | 19 | 0.02* | 0.12 |
| | Lateral_Right - Upper_Right | -3.02 | 19 | 0.01** | 0.12 |
| | Lateral_Right - Lower_Left | -1.10 | 19 | 0.29 | 0.45 |
| | Lateral_Right - Lower_Right | -1.97 | 19 | 0.06 | 0.19 |
| | Lateral_Right - Center | -0.09 | 17 | 0.93 | 0.97 |
| | Upper_Left - Upper_Right | -2.01 | 21 | 0.06 | 0.19 |
| | Upper_Left - Lower_Left | -1.52 | 20 | 0.14 | 0.38 |
| | Upper_Left - Lower_Right | -2.20 | 20 | 0.04* | 0.19 |
| | Upper_Left - Center | -1.22 | 18 | 0.24 | 0.43 |
| | Upper_Right - Lower_Left | -1.28 | 20 | 0.21 | 0.43 |
| | Upper_Right - Lower_Right | -0.93 | 19 | 0.36 | 0.51 |
| | Upper_Right - Center | -1.21 | 18 | 0.24 | 0.43 |
| | Lower_Left - Lower_Right | -1.97 | 19 | 0.06 | 0.19 |
| | Lower_Left - Center | -0.50 | 17 | 0.63 | 0.77 |
| | Lower_Right - Center | -0.62 | 17 | 0.54 | 0.71 |
| poststim - prestim | Lateral_Left - Lateral_Right | -2.52 | 6 | 0.05* | 0.06 |
| | Lateral_Left - Upper_Left | -2.19 | 6 | 0.07 | 0.08 |
| | Lateral_Left - Upper_Right | -3.91 | 6 | 0.01** | 0.02* |
| | Lateral_Left - Lower_Left | -6.59 | 6 | 0.00*** | 0.01** |
| | Lateral_Left - Lower_Right | -3.46 | 6 | 0.01* | 0.03* |
| | Lateral_Left - Center | -2.02 | 4 | 0.11 | 0.11 |
| | Lateral_Right - Upper_Left | -2.83 | 19 | 0.01* | 0.03* |
| | Lateral_Right - Upper_Right | -3.69 | 20 | 0.00** | 0.01** |

| | | | | |
|---|---|---|---|---|
| Lateral_Right - Lower_Left | -3.98 | 19 | 0.00*** | 0.01** |
| Lateral_Right - Lower_Right | -2.55 | 19 | 0.02* | 0.03* |
| Lateral_Right - Center | -2.23 | 17 | 0.04* | 0.05 |
| Upper_Left - Upper_Right | -3.57 | 20 | 0.00** | 0.01** |
| Upper_Left - Lower_Left | -2.48 | 19 | 0.02* | 0.03* |
| Upper_Left - Lower_Right | -1.95 | 19 | 0.07 | 0.08 |
| Upper_Left - Center | -3.46 | 18 | 0.00** | 0.01** |
| Upper_Right - Lower_Left | -3.47 | 18 | 0.00** | 0.01** |
| Upper_Right - Lower_Right | -2.25 | 18 | 0.04* | 0.05 |
| Upper_Right - Center | -1.79 | 17 | 0.09 | 0.10 |
| Lower_Left - Lower_Right | -2.83 | 18 | 0.01* | 0.03* |
| Lower_Left - Center | -2.59 | 17 | 0.02* | 0.03* |
| Lower_Right - Center | -2.79 | 17 | 0.01* | 0.03* |

*Note*. df = degrees of freedom. * $p < .05$. ** $p < .01$. *** $p < .001$
